# JAK Inhibitors Suppress Colon Cancer Cachexia-Associated Anorexia and Adipose Wasting in Mice

**DOI:** 10.1101/2020.01.28.923391

**Authors:** Gurpreet Arora, Arun Gupta, Tong Guo, Aakash Gandhi, Aaron Laine, Chul Ahn, Dorothy Williams, Puneeth Iyengar, Rodney Infante

## Abstract

**Background:** Cachexia (CX), a syndrome of muscle atrophy, adipose loss, and anorexia, is associated with reduced survival in cancer patients. The colon adenocarcinoma C26c20 cell line secretes the cytokine leukemia inhibitor factor (LIF) which induces CX. We characterized how LIF promotes CX-associated weight loss and anorexia in mice through JAK-dependent changes in adipose and hypothalamic tissues.

**Methods:** CX was induced *in vivo* with C26c20 colon adenocarcinoma cells or recombinant LIF administration in the absence or presence of JAK inhibitors. Blood, adipose, and hypothalamic tissues were collected and processed for cyto/adipokine ELISAs, immunoblot analysis, and quantitative RT-PCR. CX was induced *in vitro* by stimulating differentiated adipocytes with recombinant LIF or IL-6 in the absence or presence of lipase or JAK inhibitors. These activated adipocytes were processed for lipolysis, immunoblot analysis, and RT-PCR.

**Results:** Tumor-secreted LIF induced changes in adipose tissue expression and serum levels of IL-6 and leptin in a JAK-dependent manner influencing CX-associated adipose wasting and anorexia. We identified two JAK inhibitors that block cytokine-mediated adipocyte lipolysis and IL-6 induction using an *in vitro* CX lipolysis assay. JAK inhibitors administered to *in vivo* colon cancer CX mouse models led to 1) a decrease in STAT3 phosphorylation in hypothalamic and adipose tissues, 2) a reverse in the CX serum cyto/adipokine signature, 3) a delay in colon cancer CX-associated anorexia and adipose loss, and 4) an improvement in overall survival.

**Conclusions:** JAK inhibitors suppress cytokine-associated adipose loss and anorexia in multiple *in vitro* and *in vivo* models of cancer CX.

## INTRODUCTION

Cachexia (CX) represents a wasting syndrome consisting of muscle atrophy, adipose loss, and anorexia. It is observed in up to 50% of patients with solid tumors.^1^ Cancer patients with CX have greater than a 50% reduction in overall survival when compared to stage matched patients without cachexia.^1^Even when CX is identified in early stage cancer patients, survival is not improved because of a potential lack of effective therapeutic interventions.^2^ Although CX has been recognized for more than half a century, most preclinical studies and clinical trials targeting the immune system (TNF-α, IL-6), appetite stimulation, and muscle regeneration have failed to durably improve the syndrome.^3^ The majority of studies on CX have focused on the sarcopenia of this wasting syndrome, with less emphasis on adipose loss or anorexia. Recent evidence suggests that blocking adipocyte lipolysis using global lipase null mice limited not only the adipose wasting but also the sarcopenia observed in murine models of cancer CX.^4^ Additionally, Zimmers and colleagues demonstrated that pancreatic cancer cachexia patients can have adipose wasting without sarcopenia and an associated decrease in survival.^5^ Therefore, identification and complete characterization of CX factors and the common mechanisms that they utilize to induce adipose wasting and anorexia may lead to an effective treatment for patients with cancer CX.

When transplanted into syngeneic mice, the murine colon adenocarcinoma cell line C26c20 promotes CX-associated adipose wasting and anorexia.^6^ Previously, we created an *in vitro* CX screen to identify cancer-secreted molecules that can contribute to CX-associated adipose wasting. This screen identified leukemia inhibitory factor (LIF) as a CX-inducing molecule secreted from the C26c20 colon cancer line. In an analysis of more than 30 cancers in the TCGA database, LIF is most highly expressed in gastrointestinal, thoracic, and genitourinary cancers^6^ which are all associated with cachexia. Two recent papers suggested that LIF is critical to pancreatic cancer development,^7, 8^ further illustrating the importance of this molecule to CX-associated cancers. LIF is a 21 kDa protein in the IL-6 family of cytokines that binds to its receptor, LIFR-α, and the co-receptor gp130 inducing JAK-STAT signaling.^9^ Recombinant LIF (rLIF) administered to wild-type mice causes adipose and body weight loss reproducing the CX phenotype.^6^ LIF induces this wasting through JAK/STAT signaling centrally to reduce food intake and peripherally to stimulate adipocyte lipolysis beyond the anorexic effects.^6^ As rLIF induces the cachexia-associated adipose loss, there is a corresponding decrease in serum levels of leptin. Leptin is an adipokine, a cytokine-like molecule produced by adipose tissue, which regulates appetite through JAK/STAT signaling of the hypothalamus in the setting of changes in adipose levels.^10, 11^ As rLIF induces CX-associated adipose loss, the reduction of adipose leptin compensates for rLIF’s anorexic effect.

In the present study, we show that the colon adenocarcinoma C26c20 and rLIF-driven cachexia mouse models have increased serum LIF and IL-6 with decreased leptin, defining a CX signature in mice. Consistent with these serum findings, adipose mRNA levels of LIF, IL-6, and leptin from both cachexia models were similarly altered. By showing that LIF was still able to induce CX in *IL-6^−/−^* mice, we demonstrated that both LIF and IL-6 could independently promote anorexia and adipose loss. We therefore hypothesized that inhibition of the JAK signaling pathway would suppress the CX phenotype, since the molecules that are altered in the CX cyto/adipokine signature (LIF, IL-6, and leptin) all signal through this pathway.^12^ A screen of candidate JAK inhibitors in our *in vitro* CX adipocyte lipolysis assay led us to test tofacitinib and ruxolitinib *in vivo*. These JAK inhibitors are effective in the treatment of ulcerative colitis, rheumatoid arthritis, and myelofibrosis.^13-15^ The independent administration of either JAK inhibitor to cytokine- or tumor-driven models of cancer CX led to decreased STAT3 phosphorylation in adipose and hypothalamic tissues with a concomitant suppression of CX-associated changes to adipose mRNA expression and serum levels of cyto/adipokines. The net effect of these changes resulted in a mitigation of rLIF-induced anorexia and adipose/body weight loss and additionally led to an improvement of survival in the C26c20 colon cancer CX model. These studies suggest that: 1) Colon cancer-secreted LIF induces CX through signaling on the hypothalamus and adipose tissue altering levels of multiple cyto/adipokines, and 2) JAK inhibition of these signaling events suppresses CX-associated adipose loss and anorexia long enough to lead to an improvement in cancer CX survival.

## METHODS

See Supplementary Materials for a more detailed Materials and Methods.

### Mouse Studies

Male wild-type Balb/c mice and C57BL/6J were obtained from Charles River Laboratories or Jackson Laboratories at approximately 8 weeks of age. *IL-6^−/−^* mice (B6.129S2-IL6^tm1Kopf^/J, 002650) were obtained from Jackson Laboratories at approximately 7 weeks of age. All mice were allowed to acclimate in UT Southwestern animal facilities before experimentation for at least 1 week. Animals were kept in a temperature-controlled facility with a 12 h light/dark cycle and were fed normal chow diet and provided water ad libitum. Approximately 100 g of standard chow diet was placed in each cage. When food reached approximately 50 g per cage, it was replenished to approximately 100 g. Food was weighed at the same time daily and compared to the previous day’s weight to calculate the 24 h food intake per cage. Body weight was measured using a standard balance (digital Soenhle scale). Adipose tissue mass and lean tissue mass were measured longitudinally using ECHO MRI (ECHO Medical Systems) at 9AM at the indicated time points. Whole blood was drawn from tail vein bleed longitudinally or by cardiac puncture at terminal time points. Serum was obtained by subjecting the whole blood to centrifugation at 960 × g at 4 °C for 10 min. Supernatant was removed followed by protein concentration quantification using a bicinchonianic acid kit (Pierce). For analysis of serum cytokine changes, 25 μl of mouse serum was diluted with 25 μl of PBS and sent to Eve Technologies (Calgary, Canada) for ELISA analysis of multiple cytokines. For analysis of serum leptin or IL-6, serum dilution and ELISA analysis was performed as performed per kit directions from Crystal Chem and Sigma, respectively. The rest of the serum was stored at −80 °C for future blood analysis. For tumor studies, 0.75-1 × 10^7^ C26c20 cells in 100 μl PBS were injected into the right hind flank of mice at day 0, and tumor volume was calculated by taking half of the product of the caliper (VWR) measurements of length, width and breath at the indicated time points. At the end of the experiments, mice were euthanized at the indicated time point in non-tumor experiments or within 12 ^hours of expected death in tumor studies as recommended by IACUC using a CO^2 ^chamber and organs^ were collected and snap frozen.

### Adipocyte Lipolysis Assay

Media glycerol concentration from differentiated adipocytes was measured for each condition in triplicate as previously described.^6^

### Statistical Analysis

Data is presented as mean ± SEM, dot plots ± SEM, or dot plots with bars representing mean ± SEM. A Student’s *t* test was used to determine differences between groups at distinct time points. A Generalized Estimating Equation approach was used to determine differences between groups over time. Kaplan Meier analysis was conducted for survival with statistical evaluation using the Gehan-Breslow-Wilcoxon approach. For some animal studies, the ROUT method was used to remove outliers followed by ANOVA with a multiple comparison post-test (Dunnett’s or Tukey’s). Significance was considered if *p* < 0.05.

### Study Approval

All animal studies were conducted under an Institutional Animal Care and Use Committee approved protocol at UT Southwestern Medical Center (Dallas, Texas).

## RESULTS

### Serum and Adipose mRNA Cyto/Adipokine Levels in the C26c20 CX Mouse Model

To verify that the increased LIF and decreased leptin serum levels observed previously in the rLIF-induced CX model^6^ were also altered similarly in the colon cancer CX model, we evaluated the cytokine and leptin levels in serum from the C26c20 CX mouse model. Serum was collected from these colon cancer-bearing mice for cyto/adipokine ELISA analysis after each animal lost 30-40% of its adipose mass. These mice had a significant serum increase in LIF (Figure 1A) and a decrease in leptin (Figure 1B), similar to findings from the rLIF-induced CX model.^6^ In addition to the expected changes in LIF and leptin levels in the C26c20 colon cancer CX model, we also observed a 10-fold increase in serum IL-6 levels. Other groups have demonstrated an association between altered serum IL-6 levels and CX.^16-18^ Furthermore, we generalized these findings of increased serum IL-6 and LIF across multiple syngeneic murine cancer cachexia models, including those created with LLC and 4T1 tumor cells (data not shown).

**Figure 1.**
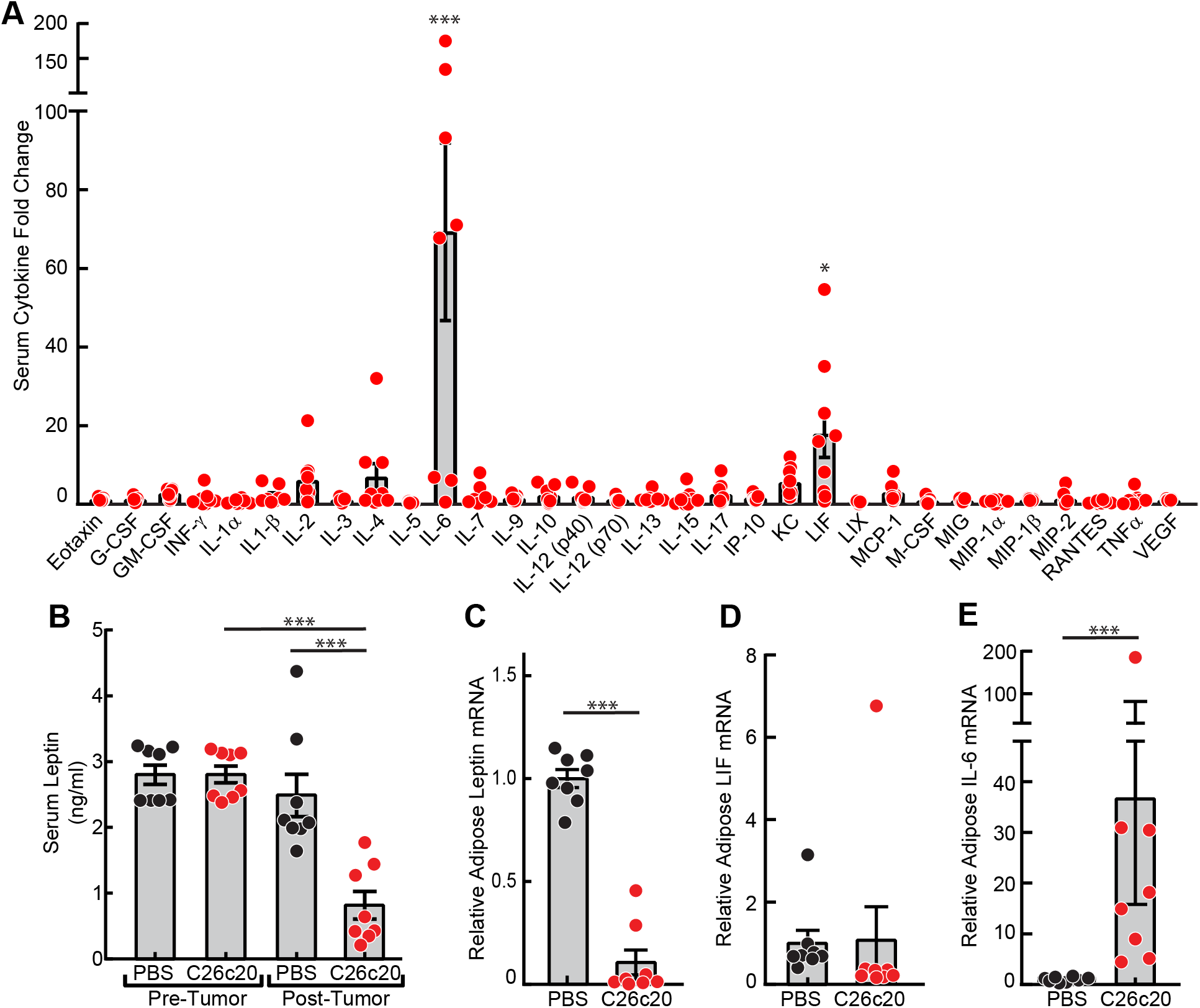
Serum and Adipose mRNA Levels of Cyto/Adipokines in a Colon Cancer Cachexia Mouse Model. Chow-fed Balb/c mice (10-week-old males) were injected s.c. in the right flank with PBS in the absence or presence of C26c20 cells. A-B) Serum Cyto/Adipokine levels. Blood was collected by tail vein bleed on day 0 and by cardiac puncture after each mouse lost 30-50% of its adipose mass as measured by ECHO MRI. Serum was isolated and the cytokine and adipokine levels were measured as described in *Methods*. C-E) Adipose cyto/adipokine mRNA expression levels. Epididymal white adipose tissue was harvested at sacrifice for measurement of the indicated mRNA by quantitative RT-PCR. For each gene, the amount of mRNA from PBS-treated mice is set to 1, and mRNA amounts from C26c20-bearing mouse adipose tissue are expressed relative to this reference value. The average C_t_ values for β-actin (invariant control) for PBS- and C26c20-administered mice were 18.0 and 18.7, respectively. The average PBS-administered mice C_t_ values for leptin, LIF, and IL-6 were 22.1, 28.2, and 33.1, respectively. Data is shown as dot plots with bars representing mean ± SEM (A-E) of five (A) or eight (B-E) mice. *p<0.05 and ***p<0.001 based on use of a ROUT method (Q=0.001) to remove outliers followed by an ANOVA and Dunnett’s multiple comparison post-test comparing the relative change in the indicated serum cytokine relative to VEGF (A) or to PBS control (B-E). These results were confirmed in at least two independent experiments.

Considering that adipose is the primary source of leptin,^19^ we next tested whether the cancer-induced changes in serum cyto/adipokines paralleled the changes in adipose mRNA expression levels. As expected, mice bearing C26c20 tumors demonstrated a decrease in mRNA expression of leptin in their white adipose tissue (WAT) compared to mice receiving PBS alone (Figure 1C). Although WAT from tumor-bearing mice did not exhibit an increase in their LIF mRNA expression (Figure 1D), there was a ~ 10-fold increase in the mRNA expression of IL-6 in WAT from tumor-bearing mice compared to vehicle-treated mice (Figure 1E). This data suggests that cancer secreted factors, such as LIF, not only cause adipose wasting/lipolysis, but also change the expression profile of cyto/adipokines in this tissue amplifying the CX signal.

### Serum and Adipose mRNA Cyto/Adipokine Levels in the rLIF CX Mouse Model

Knowing that rLIF is able to alter serum leptin levels, we hypothesized that it also increases IL-6 serum levels matching changes observed in *in vivo* cancer CX models. To test this hypothesis, we evaluated serum levels and adipose mRNA expression levels of LIF, IL-6, and leptin in our rLIF-driven CX model (Figure 2). As rLIF-injected mice lost fat mass (Figure 2A), there was a parallel increase in serum LIF (Figure 2B) and IL-6 (Figure 2C) with a corresponding decrease in serum leptin (Figure 2D).

**Figure 2.**
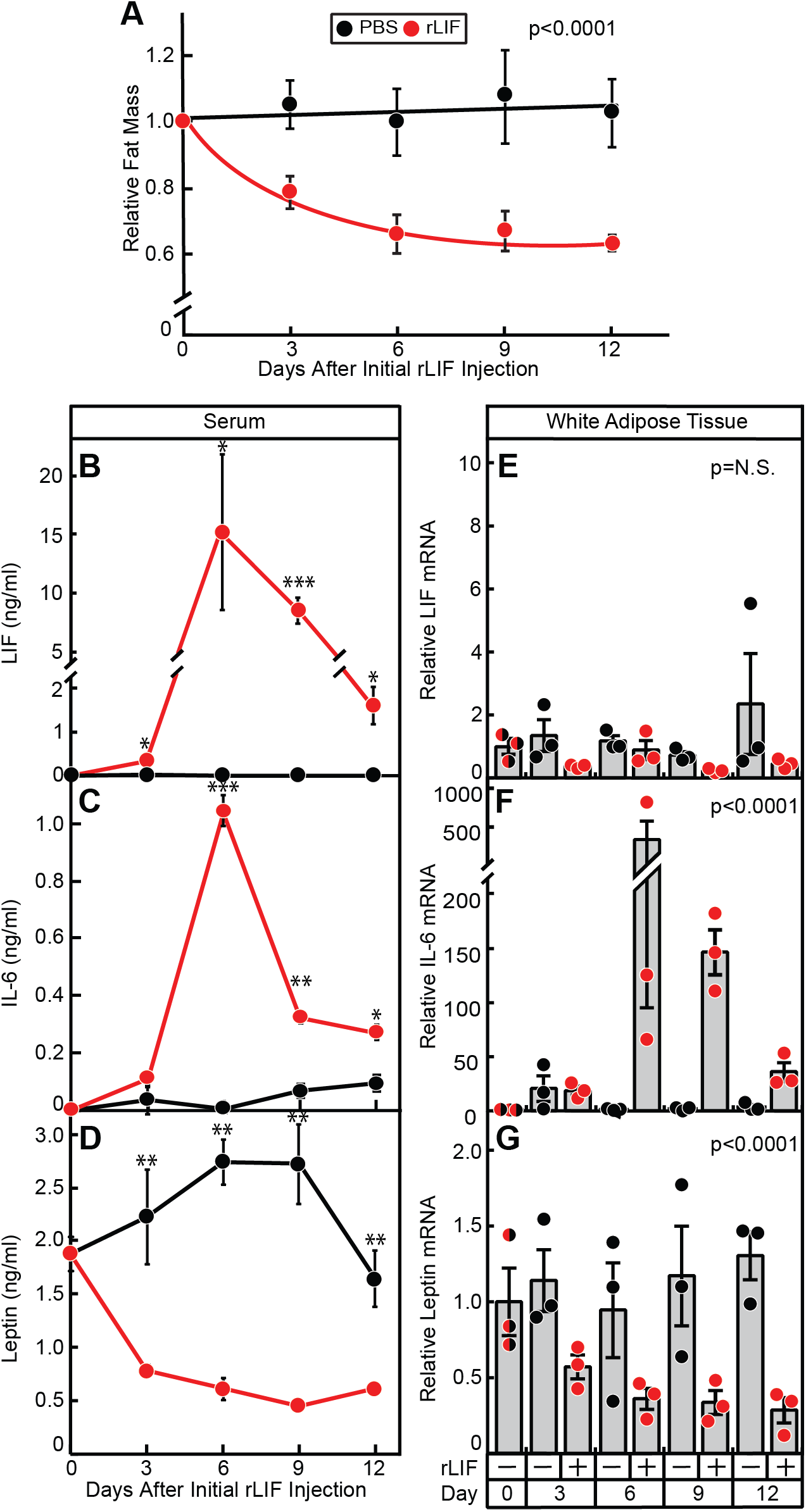
Serum and Adipose mRNA Levels of Cyto/Adipokines in the rLIF Cachexia Mouse Model. Chow-fed C57BL/6J mice (10-week-old males) were injected i.p. with PBS in the absence or presence of recombinant LIF at 80 μg/kg body weight twice daily and fat mass (A) was measured by ECHO MRI. Fat mass is shown relative to the average day 0 reference value for each respective cohort which was 3.07 and 3.16 g for the PBS- and rLIF-treated mice, respectively. B-G) Serum levels and adipose tissue mRNA expression of cyto/adipokines. Every 3 days, 3 mice from each cohort were sacrificed for harvesting of serum for cyto/adipokine ELISA (B-D) and epididymal white adipose tissue for measurement of the indicated mRNA by quantitative RT-PCR (E-G) as described in *Methods*. For each gene, the amount of mRNA from day 0 is set to 1, and mRNA amounts from adipose tissue from the indicated days are expressed relative to this reference value (E-G). The average C_t_ values for β-actin (invariant control) for day 0, 3, 6, 9, and 12 for PBS-treated mice were 18.8, 18.8, 18.9, 18.5, and 18.7, respectively, and for rLIF-treated mice were 18.8, 18.0, 18.5, 17.9, and 17.8, respectively. The average day 0 C_t_ values for LIF, IL-6, and leptin were 26.4, 33.2, and 21.8, respectively. Data is shown as mean ± SEM (A-D) or dot plots with bars representing mean ± SEM (E-G) of three mice. *p<0.05, **p<0.01, and ***p<0.001 based on use of Student’s *t*-test (B-D) or Generalized Estimating Equation approach comparing the two groups over time with rLIF-treated mice as the reference value (A, E-G). These results were confirmed in at least two independent experiments.

To determine whether adipose mRNA expression changes correlated with changes in serum cyto/adipokine levels induced by LIF, we collected RNA from WAT of rLIF- and vehicle-treated mice for quantitative RT-PCR analysis. Adipose tissue from mice treated with rLIF had a greater than 50-fold increase in IL-6 mRNA expression (Figure 2F) and an ~5-fold decrease in leptin mRNA expression (Figure 2G) with no significant change in LIF expression (Figure 2E). The changes of IL-6 and leptin adipose mRNA expression correlated with the changes observed in their respective serum levels (compare Figures 2C and 2F, 2D and 2G). Overall, the simple model of rLIF-induced CX had a similar serum and adipose signature to that of the complex C26c20 colon cancer CX model (see Figure 1) with a net result of increased serum LIF and IL-6, with a corresponding decrease in leptin.

### Evaluation of rLIF-induced CX in an *IL-6*^−/−^ Mouse Model

As described previously, rLIF injected into C57BL/6J mice causes a ~5-10% muscle loss and an ~30-40% adipose loss resulting in an ~10-15% body weight loss mimicking a CX phenotype.^6^ IL-6 has also been reported to induce CX *in vivo*.^16-18^ We have shown that LIF treatment leads to an increase in serum IL-6 (Figure 2C) and adipose IL-6 expression (Figure 2F) *in vivo.* Therefore, we next determined whether rLIF-associated CX is dependent on its induction of IL-6. PBS in the absence or presence of rLIF was injected into *IL-6^+/+^* or *IL-6^−/−^* mice and changes in food intake, body weight, and fat and lean mass by ECHO MRI were measured over time (Figure 3). Figure 3A shows that circulating concentrations of LIF were elevated in both *IL-6^+/+^* and *IL-6^−/−^* mice administered rLIF at day 21 compared to day 0. As expected, circulating levels of IL-6 were also increased with rLIF administration in *IL-6^+/+^* mice, but not in *IL-6^−/−^* or PBS-treated *IL-6^+/+^* mice at day 21 (Figure 3B). Mice receiving rLIF had reduced food intake in both *IL-6^+/+^* and *IL-6^−/−^* mice compared to mice receiving PBS during the first 9 days of the experiment (Figure 3C). Both *IL-6^+/+^* and *IL-6^−/−^* mice also demonstrated decreased fat mass (Figure 3D) and body weight (Figure 3E) when treated with recombinant rLIF compared to control conditions treated with PBS. Although an increase in IL-6 will promote a CX phenotype, LIF’s induction of CX was not dependent on its increase in serum IL-6. Therefore, both molecules are likely driving the CX phenotype in cancer models.

**Figure 3.**
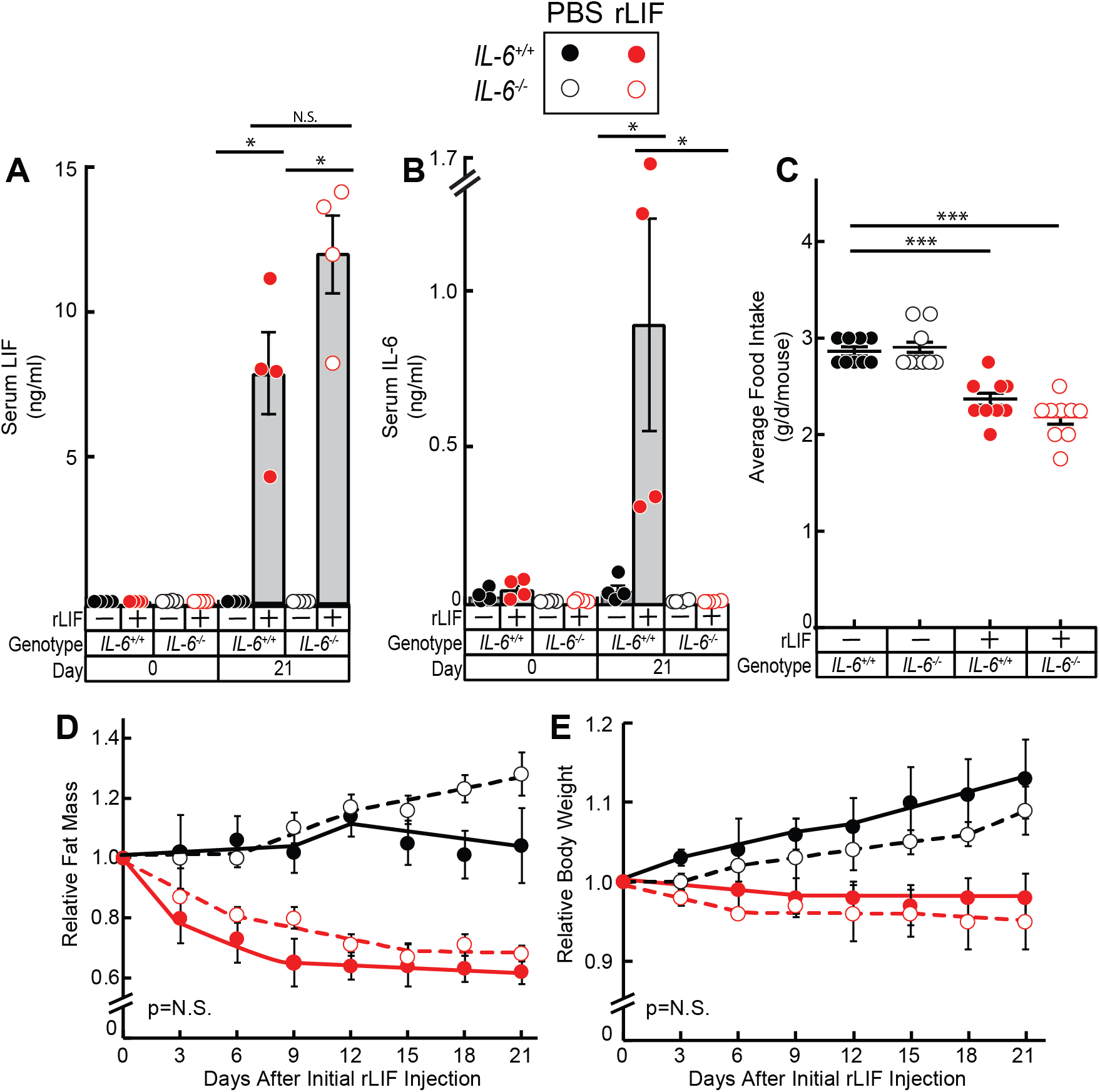
LIF Induces Cachexia-Associated Anorexia and Adipose/Body Weight Loss in *IL-6*^−/−^ mice. Chow-fed *IL-6^+/+^* and *IL-6^−/−^* C57BL/6J mice (8-week-old males) were injected i.p. with PBS in the absence or presence of rLIF at 80 μg/kg body weight twice daily. Serum was collected on day 0 and day 21 for evaluation of LIF (A) and IL-6 (B) levels by ELISA as described in *Methods*. Food intake (C), ECHO MRI measurements of fat mass (D), and body weight (E) were measured at the indicated time points. Fat mass and body weight are shown relative to the average day 0 reference value for each respective cohort. The average values for fat mass at day 0 of the PBS- and rLIF-treated IL-6^+/+^ mice were 2.8 and 2.4 g, respectively, and for the IL-6^−/−^ mice were 3.6 and 3.5 g, respectively. The average values for body weight at day 0 of the PBS- and rLIF-treated IL-6^+/+^ mice were 23 and 24 g, respectively, and for the IL-6^−/−^ mice were 24 and 24 g, respectively. Data is shown as dot plots with mean ± SEM (A-C) or each value represents mean ± SEM (D-E) of four mice. ^*^p<0.05 and ^***^p<0.001 based on Student’s *t*-test (A-C) or based on use of Generalized Estimating Equation approach comparing IL-6^+/+^ to IL-6^−/−^ mice treated with either PBS or rLIF (D-E).

### IL-6 mRNA Expression in Cytokine-stimulated Differentiated Adipocytes

To better understand how LIF and the subsequent increase in IL-6 can stimulate adipose tissue to promote CX-associated wasting, we studied the effect of these cytokines effects on *in vitro* differentiated adipocytes in relation to lipolysis and changes in mRNA expression of cyto/adipokines. As shown in Figure 4A, both wild-type rLIF and recombinant IL-6 (rIL-6) increased adipocyte lipolysis, whereas the mutant rLIF K159A had no effect. The mutant rLIF K159A is unable to stimulate lipolysis of differentiated adipocytes *in vitro* or promote CX-associated adipose wasting when administered in mice.^6^ Compared to rIL-6, rLIF stimulated lipolysis at much lower concentrations. However, rIL-6 caused ~ 3-fold higher level of maximum lipolysis than rLIF. Wild type rLIF and rIL-6, but not rLIF K159A, stimulated the phosphorylation of STAT3 in adipocytes (Figure 4B, *top immunoblot*) at similar concentrations to those necessary to stimulate lipolysis (see Figure 4A).

**Figure 4.**
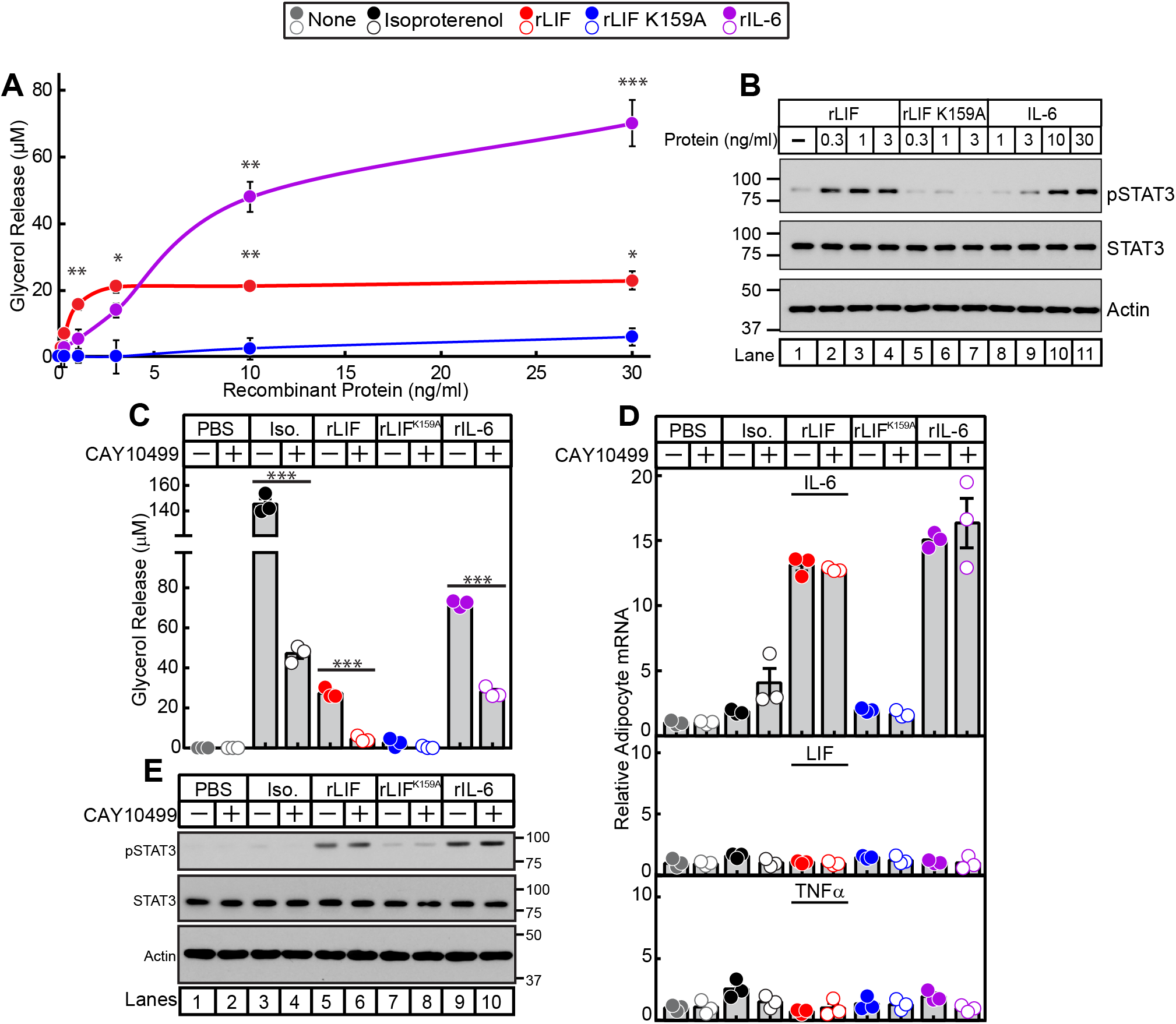
Lipolysis, JAK/STAT Activation, and mRNA Expression of Cytokine-stimulated Adipocytes. Differentiated adipocytes in a 12-well format were treated in a final volume of 1.5 ml of medium E supplemented with the indicated concentration of recombinant LIF, LIF K159A, or IL-6 (A-B) or 10 nM isoproterenol, 3 ng/ml rLIF, 3 ng/ml rLIF K159A, or 50 ng/ml rIL-6 (C-E) in the absence or presence of 10 μM CAY10499. After incubation for 20 h at 37 °C, medium was collected and glycerol concentration was measured (A,C) and cells were harvested for measurement of the indicated mRNA by quantitative RT-PCR (D) or subjected to IB analysis (B,E) with the indicated antibody as described in *Methods.* For each gene (D), the amount of mRNA from PBS-treated adipocytes in the absence or presence of CAY10499 is set to 1. The average C_t_ values for cyclophilin (invariant control) for the PBS-, isoproterenol-, rLIF-, rLIF K159A-, and IL-6-treated adipocytes in the absence of CAY10499 were 20.3, 20.9, 20.4, 20.3, and 20.7, respectively, and in the presence of CAY10499 were 20.4, 21.4, 20.6, 20.5, and 20.8, respectively. The average reference Ct values for IL-6 (*upper panel*), LIF (*middle panel*), and TNFα (*lower panel*) in PBS-treated adipocytes in the absence of CAY10499 were 28.3, 29.2, and 33.1, respectively, and in the presence of CAY10499 were 28.6, 29.2, and 33.8, respectively. Data is shown as mean ± SEM (A) or as dot plots with mean ± SEM (C,D) of three wells. *p<0.05, **p<0.01, and ^***^p<0.001 based on Student’s *t*-test comparing conditions to rLIF K159A-treated adipocytes (A) or the PBS-treated adipocytes (C-D). These results were confirmed in at least three independent experiments.

In adipocytes, triglycerides are sequentially hydrolyzed by adipose triglyceride lipase, hormone sensitive lipase (HSL), and monoacylglycerol lipase, each of which sequentially removes one fatty acid molecule to produce glycerol and fatty acids. CAY10499 is a commercially available pan inhibitor of these lipases.^20, 21^ To evaluate if cytokine-stimulated adipocyte lipolysis is required for the induction of cyto/adipokine mRNA expression, we performed quantitative RT-PCR for mRNA expression of differentiated adipocytes that were treated with either vehicle, isoproterenol, wild-type rLIF, mutant rLIF K159A, or rIL-6 in the absence or presence of lipase inhibitor CAY10499. Isoproterenol is a β-adrenergic agonist that enhances lipolysis by increasing cAMP stimulating the phosphorylation and activation of the lipase HSL.^22^ CAY10499 was able to suppress adipocyte lipolysis induced by isoproterenol, rLIF, and rIL-6 (Figure 4C). However, cytokines were still able to activate adipocytes in the presence of CAY10499 as demonstrated by cytokine-induced phosphorylation of STAT3 (Figure 4E). Although isoproterenol-treated adipocytes had a greater than 2-fold increase in lipolysis relative to rIL-6- or rLIF-treated adipocytes (Figure 4C), there was no significant increase in IL-6 mRNA expression (Figure 4D, *upper panel*). Contrarily, adipocytes treated with wild-type rLIF or rIL-6 had a greater than 10-fold increase in relative IL-6 mRNA expression, which remained elevated even in the presence of lipase inhibitor CAY10499. The mRNA expression of LIF (*middle panel*) and TNFα (*lower panel*) were unchanged in β-adrenergic-induced or cytokine-induced adipocytes in the absence or presence of CAY10499. We could not explore the cytokine regulation of leptin mRNA expression since leptin’s mRNA levels are undetectable in our *in vitro* differentiated adipocyte model. Combined, the results of these experiments demonstrated that cytokines are able to stimulate IL-6 expression in adipocytes independent of their induction of lipolysis.

### JAK Inhibition Blocks Lipolysis, JAK/STAT Activation, and IL-6 mRNA Expression in Cytokine-stimulated Differentiated Adipocytes

Having demonstrated that CX factors LIF and IL-6 are both increased in cancer CX, an ideal inhibitor would block a common pathway these cytokines activate to induce adipose loss, anorexia, and muscle wasting. Since LIF and IL-6 both induce JAK/STAT activation in adipocytes (see Figure 4B) and in the hypothalamus^6, 23^, we next screened JAK inhibitors for their ability to block *in vitro* cytokine-stimulated adipocyte lipolysis, JAK/STAT activation, and IL-6 mRNA expression. We incubated adipocytes with isoproterenol or rIL-6 in the presence of increasing concentrations of multiple JAK inhibitors including tofacitinib, ruxolitinib, decernotinib, and filgotinib (Figure 5A). Beta-adrenergic agonist isoproterenol-induced adipocyte lipolysis was not affected by any JAK inhibitor. Adipocyte lipolysis induced by rIL-6 was significantly inhibited when adipocytes were incubated with tofacitinib or ruxolitinib.

**Figure 5.**
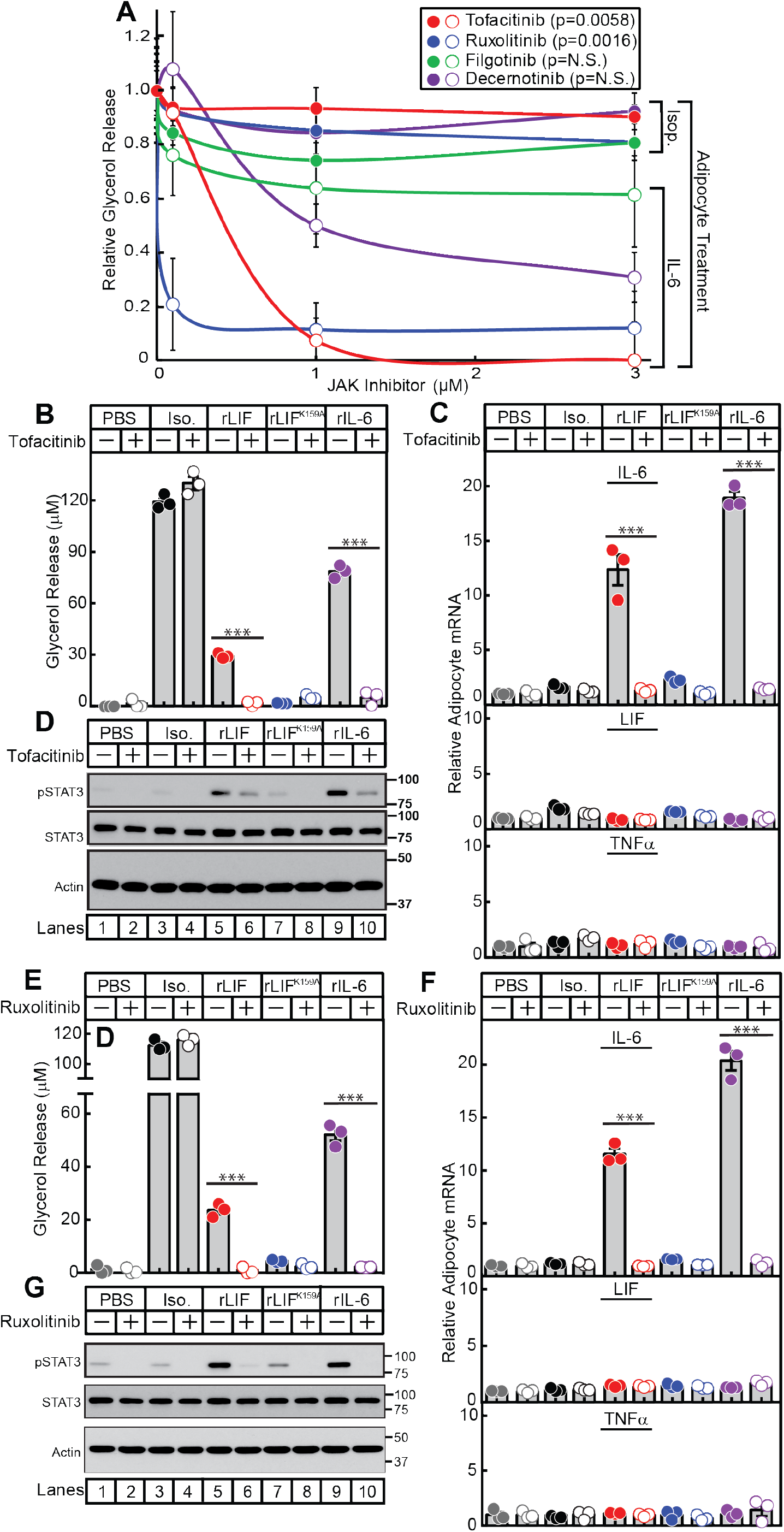
Cytokine Stimulation of Adipocytes in the Presence of Janus Kinase Inhibitors. Differentiated adipocytes in a 12-well format were treated in a final volume of 1.5 ml of medium E with 10 nM isoproterenol, 50 ng/ml rIL-6, or 3 ng/ml rLIF in the absence or presence of the indicated concentration of JAK inhibitor (A), 10 μM tofacitinib (B-C), or 3 μM ruxolitinib (E-F). After incubation for 20 h at 37 °C, medium was collected and glycerol concentration was measured (A,B,E) and cells were harvested for measurement of the indicated mRNA by quantitative RT-PCR (C,F) or subjected to IB analysis (D,G) with the indicated antibody as described in *Methods.* For each gene (C,F), the amount of mRNA from PBS-treated adipocytes in the absence or presence of the JAK inhibitor are set to 1. The average C_t_ values for cyclophilin (invariant control) for the PBS-, isoproterenol-, rLIF-, rLIF K159A-, and IL-6-treated adipocytes in the absence of tofacitinib were 20.3, 20.7, 20.4, 20.3, and 20.4, respectively, and in the presence of tofacitinib were 20.5, 20.9, 20.6, 20.6, and 20.6, respectively. The average reference Ct values for IL-6 (*upper panel*), LIF (*middle panel*), and TNFα (*lower panel*) in PBS-treated adipocytes in the absence of tofacitinib were 28.5, 29.1, and 33.5, respectively, and in the presence of tofacitinib were 30.2, 29.2, and 33.7, respectively. F) The average C_t_ values for cyclophilin (invariant control) for the PBS-, isoproterenol-, rLIF-, rLIF K159A-, and IL-6-treated adipocytes in the absence of ruxolitinib were 20.5, 20.9, 20.5, 20.6, and 20.8, respectively, and in the presence of ruxolitinib were 20.8, 21.1, 20.9, 20.9, and 21.2, respectively. The average reference Ct values for IL-6 (*upper panel*), LIF (*middle panel*), and TNFα (*lower panel*) in PBS-treated adipocytes in the absence of ruxolitinib were 28.4, 28.5, and 32.7, respectively, and in the presence of ruxolitinib were 30.2, 29.1, and 33.5, respectively. Data is shown as mean ± SEM (A) or as dot plots with mean ± SEM (B-C,E-F) of three wells. ^***^p<0.001 based on Student’s *t*-test comparing cytokine-treated adipocytes to isoproterenol-treated adipocytes at 1 μM (A) or comparing conditions to the PBS-treated adipocytes (B-C,E-F). These results were confirmed in at least three independent experiments.

To evaluate whether cytokine-induced adipocyte IL-6 expression was dependent on JAK-STAT activation, we incubated adipocytes with PBS, isoproterenol, rLIF, mutant rLIF K159A, or rIL-6 in the absence or presence of tofacitinib or ruxolitinib. As shown in Figure 5B, tofacitinib significantly suppressed rIL-6 and rLIF-mediated, but not isoproterenol-mediated adipocyte lipolysis. To verify that tofacitinib inhibited JAK-mediated STAT activation, immunoblot analysis of adipocyte lysates demonstrated reduced cytokine-mediated STAT3 phosphorylation in the presence of tofacitinib (Figure 5D, top immunoblot). Tofacitinib also completely suppressed cytokine-mediated induction of adipocyte IL-6 expression (Figure 5C, *top panel*). Adipocyte mRNA expression of other cytokines, LIF (*middle panel*) and TNFα (*bottom panel*), were unchanged in adipocytes treated with PBS, isoproterenol, or cytokines in the absence or presence of tofacitinib. Another JAK inhibitor, ruxolitinib, also suppressed cytokine-mediated lipolysis (Figure 5E), phosphorylation of STAT3 (Figure 5G), and IL-6 mRNA expression (Figure 5F, top panel).

### JAK Inhibition Suppresses Anorexia and Adipose Loss in the rLIF CX Mouse Model

To demonstrate that JAK inhibition blocks anorexia and adipose wasting in the rLIF-induced CX model, we injected mice with PBS or rLIF in the absence or presence of tofacitinib (Figures 6A-D) or ruxolitinib (Figures 6E-J). The addition of tofacitinib or ruxolitinib to rLIF-treated mice restored food intake (Figure 6A and 6E), body weight (Figure 6B and 6F), and fat mass (Figure 6C and 6G) towards levels observed in vehicle-treated mice. There was no change in lean mass among all cohorts (Figure 6D and 6H) within the limited time frame of the experiment. To confirm that JAK inhibition suppressed STAT activation in target tissues relevant to appetite and wasting, STAT3 phosphorylation was measured in hypothalamic and adipose tissues, respectively, by immunoblot analysis. Mice treated with rLIF and ruxolitinib (*lanes 10-12*) had decreased STAT3 phosphorylation compared to rLIF-treated mice in the absence of ruxolitinib (*lanes 7-9*) in hypothalamic (Figure 6I) and adipose (Figure 6J) tissues.

**Figure 6.**
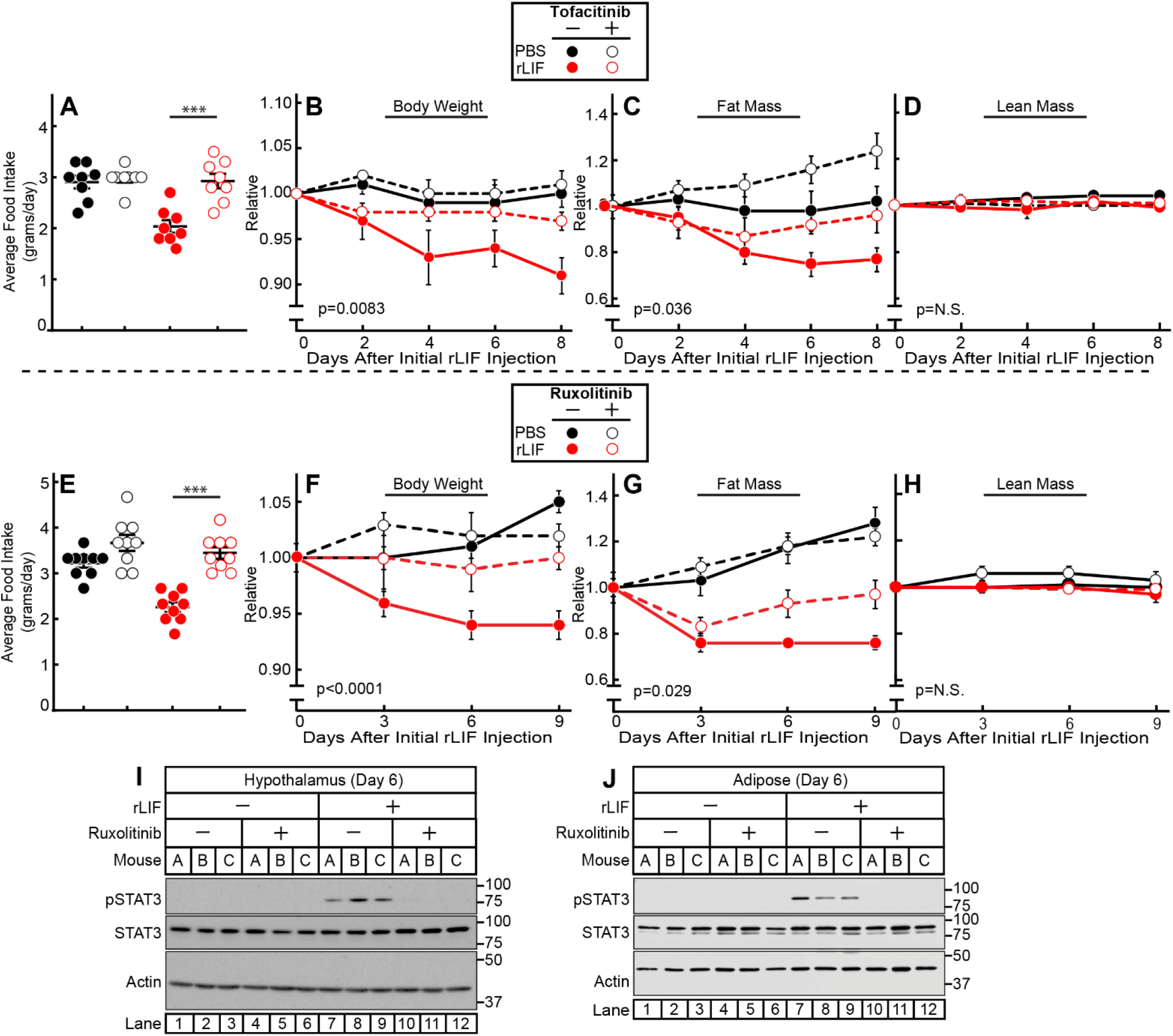
JAK Inhibition Suppresses rLIF-mediated Cachexia. Chow-fed C57BL/6J mice (10-week-old males) were were injected i.p. with PBS in the absence or presence of rLIF at 80 μg/kg body weight twice daily. Ninety minutes after each rLIF injection, mice also received either oral gavage of 200 μl PBS containing 0.5% (*v/v*) methylcellulose and 0.1% (v/v) Tween 20 in the absence or presence of 25 mg/kg tofacitinib (A-D) or i.p. of 150 μl PBS containing 2% DMSO (*v/v*) and 30% PEG300 (*v/v*) in the absence or presence of 25 mg/kg ruxolitinib (E-J) twice daily. Food intake (A,E), body weight (B,F), ECHO MRI measurement of fat mass (C,G) and lean mass (D,H), or IB analysis of harvested hypothalamic (I) or adipose (K) tissue were measured at the indicated time points. Body weight, fat mass, and lean mass are shown relative to the average day 0 reference value for each respective cohort. The average values for day 0 for the PBS with vehicle, PBS with tofacitinib, rLIF with vehicle, and rLIF with tofacitinib were as follows: body weight (27, 25.4, 26.7, and 26.8 g), fat mass (3.6, 3.4, 3.7, and 3.6 g) and lean mass (19, 18, 18.7, and 19 g), respectively. The average values for day 0 for the PBS with vehicle, PBS with ruxolitinib, rLIF with vehicle, and rLIF with ruxolitinib were as follows: body weight (26.5, 28, 26.4, and 27 g), fat mass (2.7, 3.0, 3.0, and 2.8 g) and lean mass (20, 21, 20, and 21 g), respectively. Data is shown as dot plots with mean ± SEM (A and E) or each value represents the mean ± SEM (B-D,F-H) of four (A-D) or three (E-H) mice. These results were confirmed in at least three independent experiments.

### JAK Inhibition Suppresses Anorexia and Adipose Loss Improving Survival in the C26c20 CX Mouse Model

To test if JAK/STAT signaling pathways are an integral component of colon cancer CX, we tested JAK inhibitors in the *in vivo* C26c20 cancer CX model. Mice were injected with PBS or C26c20 cells on day 0 in the absence or presence of ruxolitinib (Figure 7). Mice implanted with colon cancer C26c20 cells and treated with ruxolitinib had an ~20-30% increase in median survival compared to mice receiving C26c20 in the absence of ruxolitinib (Figure 7A), despite the absence of any significant ruxolitinib effect on tumor growth (Figure 7B). Mice receiving both C26c20 cells and ruxolitinib had a reduction in the CX-associated-adipose loss at intermediate time points (Figure 7D, *right panel*), coinciding with decreased adipose tissue STAT3 phosphorylation (Figure 7F, *top blot, compare lanes 4 and 5*). At the time that animals were sacrificed due to CX morbidity, there was no longer a significant difference in adipose mass either in the absence or presence of ruxolitinib (Figure 7D, *right panel*). This end point coincided with increasing adipose tissue STAT3 phosphorylation even in the ruxolitinib-treated colon cancer-bearing mice (Figure 7F, top blot, *compare lanes 9,10*). Mice receiving both C26c20 cells and ruxolitinib also had a reduction in the CX-associated anorexia between days 5 and 8 (Figure 7C, *middle panel*), coinciding with decreased hypothalamic tissue STAT3 phosphorylation (Figure 7F, 4^th^ blot, *compare lanes 4 and 5*). However, these mice reached the same levels of anorexia before sacrifice (days 9-12) as the colon cancer-bearing mouse cohort receiving vehicle (Figure 7C, *right panel*), coinciding with increasing adipose tissue STAT3 phosphorylation even in the ruxolitinib-treated colon cancer-bearing mice (Figure 7F, 4^th^ blot, *compare lanes 9,10*). During intermediate time points when JAK inhibition was effective at blocking STAT3 phosphorylation in adipose tissue, there was also a significant suppression of CX-associated changes in circulating leptin (Figure 7G) and IL-6 (Figure 7H). As STAT3 phosphorylation returned in adipose tissue even in the presence of JAK inhibition, the serum levels of IL-6 and leptin approached those observed in a CX mouse without JAK inhibition.

**Figure 7.**
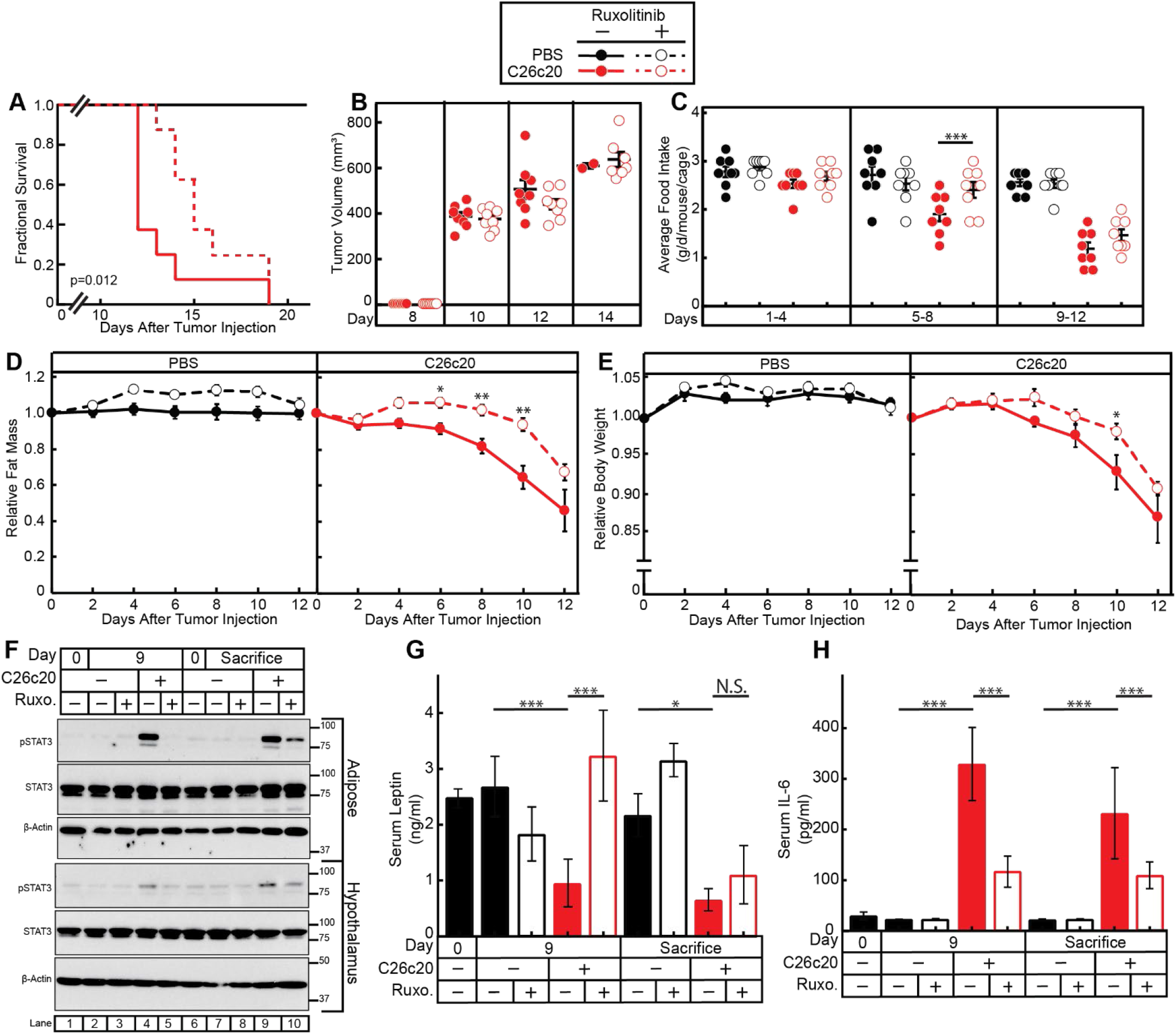
JAK Inhibition Suppresses Colon Cancer Cachexia. Chow-fed Balb/c mice (10-week-old males) were injected s.c. in right flank with PBS in the absence or presence of C26c20 cells on day 0 followed by i.p. administration of 150 μl PBS containing 2% DMSO (*v/v*) and 30% PEG300 (*v/v*) in the absence or presence of 25 mg/kg ruxolitinib twice daily thereafter. Survival (A), tumor volume (B), food intake (C), ECHO MRI of fat mass (D), body weight (E), IB analysis of harvested hypothalamic and adipose tissue with the indicated antibodies (F), and ELISA of the serum of the indicated cyto/adipokine (G-H) were measured at the indicated time points as described in *Methods*. Fat mass and body weight are shown relative to the average day 0 reference value for each respective cohort. The average values for day 0 for the PBS with vehicle, PBS with ruxolitinib, C26c20 with vehicle, and C26c20 with ruxolitinib were 3.3, 2.8, 3.2, and 3.3 g for fat mass and 24, 23, 23, and 23 g for body weight, respectively. Data is shown as Kaplan-Meier survival curve (A), dot plots with mean ± SEM (B-C, G-H), or each value represents the mean ± SEM (D-E) of eight mice (A-G) or four mice (G-H). The Gehan-Breslow-Wilcoxon approach was used to compare survival curves (A) of C26c20-bearing mice in the absence or presence of ruxolitinib. ^*^p<0.05,^**^p<0.01, and ^***^p<0.001 based on Student’s *t*-test comparing the absence to the presence of ruxolitinib in PBS- and C26c20-bearing mice (B-E) or ANOVA and Tukey’s multiple comparison post-test (G-H). These results were confirmed in at least three independent experiments.

## DISCUSSION

LIF is a tumor-secreted factor that induces CX-associated anorexia and adipose loss through actions on the hypothalamus and increased lipolysis in adipose tissue.^6^ In this study, we demonstrated that the murine colon cancer C26c20 model has a serum cyto/adipokine signature of increased LIF and IL-6 and decreased leptin, generalizable across multiple *in vivo* CX tumor models. Consistent with the serum findings, there was also an increase in IL-6 and a decrease in leptin mRNA expression in adipose tissue of cancer bearing mice. These cyto/adipokine changes in serum and adipose mRNA expression levels were consistently altered in both the C26c20 colon cancer CX model and the rLIF-administered CX model. We next considered whether LIF’s induction of CX was a consequence of its direct effect on target tissues or due to the upregulation of the other CX factor, IL-6. LIF’s ability to induce CX *in vivo* was not dependent on IL-6 since it still promoted CX-associated anorexia and wasting in *IL-6^−/−^* mice. However, this LIF-induced systemic increase of IL-6 likely enhanced LIF’s overall contribution to CX development, with IL-6 able to stimulate adipocyte lipolysis (Figure 4A), alter appetite through hypothalamic signaling,^23^ and promote the further amplification of systemic IL-6 expression (Figure 4C). The cytokine-mediated induction of IL-6 mRNA expression was also observed *in vitro* when differentiated adipocytes were stimulated by cytokines. We used this model to show that cytokine-mediated IL-6 mRNA induction was not dependent on lipolysis, but rather on JAK-STAT pathway activation. Use of inhibitors of JAK, a common signaling pathway of LIF and IL-6, suppressed CX development in rLIF-treated mice and C26c20 colon cancer-bearing mice, resulting in decreased CX-associated anorexia (Figure 6A, 6E, and 7C) and adipose loss (Figures 6C, 6G, and 7D) that correlated with a parallel decrease in STAT3 phosphorylation of the hypothalamus (Figures 6I and 7F) and adipose tissue (Figures 6J and 7F). Furthermore, JAK inhibition in these CX models normalized cytokine-driven alterations in cyto/adipokine serum levels (Figures 7G and 7H) during time points of effective inhibition of STAT3 phosphorylation of the adipose tissue (Figure 7F). Inhibiting the JAK-STAT pathway in target tissues resulted in an approximate 20-30% increase in median survival (Figure 7A), importantly, without significant change in primary tumor size (Figure 7B).

Targeting single molecules such as TNFα, IL-6, or ghrelin have not met the threshold for becoming standard of care treatments for CX.^24-27^ The lack of a durable therapy could be due to the following: 1) the previously targeted molecules are not relevant to all types of CX; 2) each patient’s CX may be driven by a unique set of factors; and/or 3) there are multiple factors upregulated in CX that contribute to the cachexia phenotype. Our results favor the third hypothesis. They suggest how a single cytokine amplifies its own signal through targeting of multiple tissues including the hypothalamus and adipose, changing circulating levels of multiple cytokines and adipokines that work synergistically to promote the CX-associated appetite and body composition changes. We hypothesize that colon cancer CX-associated tumors and/or chronic inflammation exploit an intrinsic signaling axis between the immune system, adipose and hypothalamus to enhance anorexia and wasting, reducing survival in cancer models. Tumor- or immune-secreted cytokines, such as LIF, act on the adipose tissue to promote lipolysis and alter the release of appetite regulating molecules IL-6 and leptin. LIF can also induce expression of IL-6 in other cell types, such as fibroblasts^28^ and myotubules.^29^ These altered levels of cyto/adipokines also act directly on the hypothalamus to regulate appetite. Additionally, others have shown that IL-6 can also cause CX-associated muscle atrophy in a STAT3 dependent manner.^30^ We are encouraged that a simplified rLIF-injected mouse model is appropriate to study CX since a recent publication showed that genetic silencing of *LIF* from the C26 parental tumor line led to an anticipated decrease in systemic levels of IL-6 with suppression of the CX phenotype.^31^ Therefore, our studies support the premise that CX patients have multiple circulating CX-inducing factors at any given time explaining why therapeutic interventions targeting a single molecule have been ineffective.

Because of the heterogeneity in potential factors driving the CX phenotype, there is a need to identify the common downstream signaling pathways to elucidate targets to permit sustained responses. To block LIF and its activation of target tissues, an inhibitor would have to block not only LIF’s direct effects on the hypothalamus and adipose tissue but also its indirect effects on these tissues from changing other cyto/adipokine serum levels. Therefore, we inhibited JAK since both LIF and IL-6 use this pathway when signaling target tissues. With evidence that both tofacitinib and ruxolitinib could block lipolysis and induction of IL-6 in our *in vitro* CX adipocyte assay, we tested these compounds in our *in vivo* CX models. Both JAK inhibitors independently blocked rLIF- and colon cancer-induced CX, with suppression of anorexia and adipose/body weight loss. In the colon cancer CX model, the JAK inhibitors improved median overall survival. The compounds were most effective in blocking CX-induced anorexia and adipose loss when they were able to suppress adipose and hypothalamic STAT3 phosphorylation. This data supports the importance of both anorexia and adipose loss to the CX phenotype. Ultimately, to enhance the therapeutics of these JAK inhibitors for the treatment of CX to permit a durable suppression of STAT3 phosphorylation in these target tissues, it will be necessary to optimize the pharmacokinetics and identify other selective inhibitors that block the subclass of JAK molecules regulating the cytokine-mediated signaling. It will also be important to evaluate the effect of these JAK inhibitors on the STAT3 negative feedback regulators including protein tyrosine phosphatases, suppressors of cytokine signaling, and protein inhibitor of activated STAT. The effect of each JAK inhibitor on tumor growth kinetics and immune modulation will also need to be evaluated.

In patients with myelofibrosis, ruxolitinib improved clinical symptoms and survival.^14^ Those who received ruxolitinib also had an ~3% weight gain compared to those receiving placebo who had ~2% weight loss. The patients receiving ruxolitinib who gained weight also had a decrease in serum IL-6 and an increase in serum leptin. The changes in serum cyto/adipokine levels of ruxolitinib-treated patients are consistent with a reversed CX cyto/adipokine serum signature. These clinical findings support the importance of cytokine signaling to human physiology and pathology.

In summary, our studies demonstrate the inherent complexity of treating patients with cancer CX, since the presence of even one CX factor can amplify its signal by altering the levels of other independently-acting CX and appetite-regulating molecules. Due to these changes of serum cyto/adipokines in cancer CX, our data offers an explanation for why agents targeting single molecules have been ineffective in curtailing CX progression. Our current findings indicate that targeting common pathways, such as JAK, of cytokine-mediated signaling in the adipose and hypothalamic tissues will suppress the cancer CX phenotype of hypophagia, muscle atrophy, adipose loss, and body weight loss, resulting in improved survival.

## ETHICAL STANDARDS

All authors certify that they comply with the ethical guidelines for authorship and publishing in the Journal of Cachexia, Sarcopenia, and Muscle: update 2017.^32^

## Abbreviations

CX: cachexia
IL-6: interleukin 6
JAK: janus kinase
LIF: leukemia inhibitory factor
rLIF: recombinant leukemia inhibitory factor
STAT3: signal transducer and activator of transcription 3

## ACKNOWLEDGEMENTS

We thank Michael Brown and Joseph Goldstein for their continued mentorship and valuable suggestions. We also thank Jay Horton and Arun Radhakrishnan for their valuable suggestions. We appreciate the laboratory of Joel Elmquist for help with hypothalamic processing. We thank Dorothy Williams for excellent technical assistance and Ijeoma Dukes and Lisa Beatty for cell culture assistance. This work was supported by the Burroughs Wellcome Fund Career Awards for Medical Scientists (1019692), the American Gastroenterological Association grant 2019AGARSA3, American Cancer Society grants RSG-15-061-01-TBE and IRG-17-174-13; and the National Institute of Health grants 5P01-HL20948, P30CA142543, and 5T32GM007062-44.

