## Supplement Materials for "JAK Inhibitors Suppress Colon Cancer Cachexia-Associated Anorexia and Adipose Wasting in Mice"

**SUPPLEMENTARY MATERIALS**

**Materials.** We obtained Precision Plus Protein Kaleidoscope Standards from BioRad, Inc; protease inhibitor cocktail set III from Calbiochem; phenylmethanesulfonyl fluoride (PMSF), 3-isobutyl-1-methylxanthine (IBMX), bovine serum albumin (BSA), dexamethasone, fetal bovine serum (FBS), free glycerol reagent, glycerol, dimethylsulfoxide (DMSO), methylcellulose, PEG300, endotoxin free phosphate buffered saline (PBS), and tofacitinib from Sigma. Ruxolitinib, filgotinib, and decernotinib were obtained from Selleckchem. Isoproterenol, insulin, and CAY10499 were obtained from Cayman Chemicals. Tween 20 was obtained from Fisher Scientific. Mouse leptin ELISA kit was obtained from Crystal Chem. Recombinant IL-6 was obtained from Sigma or Peprotech. We obtained monoclonal anti-β-Actin (catalog 3700), monoclonal anti-STAT3 (catalog 9139), and monoclonal anti-pSTAT3 (catalog 9138) from Cell Signaling Technology.

**Buffers and Culture Medium.** Buffer A contained 10 mM Tris-HCL (pH 6.8), 100 mM NaCl, 1% (w/v) SDS, 1 mM EDTA, 1 mM EGTA, phosphatase inhibitor cocktail Set I (EMD Millipore 524624) and Set II (EMD Millipore 524625), and protease inhibitor cocktail (1:1000). Medium A was DMEM high glucose (MilliporeSigma, catalog D6429). Medium B was medium A supplemented with 100 units/ml penicillin (Sigma), 100 µg/ml of streptomycin sulfate (Sigma) and 10% FBS (v/v). Medium C was Hyclone DMEM without sodium pyruvate and phenol red (SH30284.02; GE Lifesciences). Medium D was medium C supplemented with 100 units/ml penicillin, 100 µg/ml of streptomycin sulfate, and 1 mM sodium pyruvate (25-100-Cl; Corning). Medium E was medium D supplemented with 0.2% (w/v) BSA.

**Cell Culture.** Stock cultures of mouse adipocytes derived from the primary stromal vascular fraction (SVF) (generous gift from Rana Gupta) and C26c20 (generous gift from Kuniyasu Soda), were maintained in monolayer culture at 37°C in 10% (SVF) or 5% (C26c20) $CO_2$. Each cell line was propagated, aliquoted, and stored under liquid nitrogen. Aliquots of SVF and C26c20 were passaged for less than 4 weeks to minimize genomic instability. Every six months, the cell lines were tested for mycoplasma contamination using the MycoAlert Mycoplasma Detection Kit (Lonza). SVF pre-adipocytes were differentiated into adipocytes as previously described.[5]

**Expression and Purification of Recombinant LIF from *E. Coli*.** Wild-type and mutant versions of recombinant LIF plasmid construction, expression in *E. coli*, and protein purification were conducted as previously described.[5]

**Immunoblot Analysis.** Tissue from mouse studies were thawed on ice and subsequently 30-50 mg of tissue was combined with 1 ml of buffer A in 2 ml microtubes with ceramic beads (Omni International). Approximately 50- 60 mg of epididymal white adipose fat pad or hypothalamus was placed in a 2 ml microtubes with 1 ml of Buffer A and homogenized with a Bead Ruptor Elite apparatus (Omni International) for three cycles of 20 s at 4 m/s. The homogenized sample was kept on ice for 30 min, after which it was subjected to centrifugation at 4 °C for 10 min at 10,000 x g. The supernatant was collected, and for adipose tissue the upper lipid layer was avoided. Protein concentrations were measured using a bicinchoninic acid kit (Pierce).

For cultured cell experiments after indicated treatments, medium was removed from each well of the 12-well plates, wells were washed twice with 1 ml PBS followed by the addition of 150 μl of buffer A. Cells were harvested by scraping and transferring to a 1.7-ml Eppendorf tube. Cells were lysed using a 22-gauge needle followed by measurement of protein concentration by bicinchroninic acid kit.

Animal tissue and cell culture extracts were mixed with 5X loading buffer, heated at 95 °C for 10 minutes, and subjected to 8% to 12% SDS/PAGE (10 μg/lane for cell culture extracts, 15 μg/lane for adipose tissue extracts, and 30 μg/lane for hypothalamic tissue extracts). The electrophoresed proteins were transferred to nitrocellulose filters using the Bio-Rad Trans Blot Turbo system followed by IB staining with the following primary antibodies: IgG-Actin (1:1000 dilution), IgG- STAT3 (1:1000 dilution), and IgG-pSTAT3 (1:1000 dilution). Bound antibodies were visualized by chemiluminescence (Super Signal Substrate; Pierce) by using a 1:5,000 dilution (animal tissue extract) or 1:10,000 dilution (cells culture extract) of donkey anti-mouse IgG (Jackson ImmunoResearch) conjugated to horseradish peroxidase. Filters were exposed to Phoenix Blue X-Ray Film (F-BX810) for 1-300 s or scanned using an Odyssey FC Imager (Dual-Mode Imaging System, 2-min integration time) and analyzed using Image Studio version 5.0 (LI-COR).

**Quantitative real-time PCR**. For quantitative real-time PCR of differentiated adipocytes in cell culture with the indicated treatments of 12-well dishes, media was removed, and cells were washed twice with 1 ml of PBS. After addition of 200 µl of Buffer RLT (RNeasy Mini Kit, Qiagen, Germantown, MD), the cells were homogenized in their respective wells using a 22-gauge needle (BD Precision Glide Needle, Franklin Lakes, New Jersey). The lysates from consecutive wells treated the same way were combined and transferred to 1.5 ml Eppendorf tubes. Total RNA was prepared using the RNeasy Mini Kit (Qiagen) using the manufacturer's directions, and subjected to real time PCR analysis. The primer sequences from Integrated DNA Technologies used for PCR were as follows: IL-6 (Fwd: TCGTGGAAATGAGAAAAGAGTTG; Rev: AGTGCATCATCGTTGTTCATACA), LIF (Fwd: AGCCGTTTCCCAACAACGT; Rev: CCGTTGCCATGGAAAGAT), TNF$\alpha$ (Fwd: CTGAGGTCAATCTGCCCAAGTAC; Rev: CTTCACAGAHCAATGACTCCAAAG) and Cyclophilin (Fwd: TGGAGAGCACCAAGACAGACA; Rev: TGCCGGAGTCGACAATGAT).

For quantitative real-time PCR of adipose tissue from mouse studies, epididymal white adipose tissue was dissected and harvested from animals at time of sacrifice. Half of one epididymal white adipose fat pad of adipose tissue was placed in a 2 ml microtubes with 1 ml of Trizol (Tel_Test) and homogenized with a Bead Ruptor Elite apparatus for three cycles of 20 s at 4 m/s. These samples were then centrifuged at 14,000 rpm for 10 minutes. The lipid layer was removed and 200 µl chloroform (Sigma) was added to each sample. The tubes were vortexed for

30 seconds and centrifuged at 14,000 rpm for 10 minutes. The top layer was collected and combined with 400 μl of isopropanol and centrifuged at 14,000 rpm for 10 min. Supernatant was removed and the pellet was washed with 200 μl 70% ethanol, air dried, and re-dissolved in RNase-free water (Invitrogen). The samples were then subjected to real time PCR analysis. Along with mRNA expression analysis of LIF, IL-6, and actin (see primers above), we also measured β-actin and leptin mRNA expression with the following primers: β-actin (Fwd: CCGTGAAAAGATGACCCAGATC; Rev: CACAGCCTGGATGGCTACGT); Leptin (Fwd: CTCCATCTGCTGGCCTTCTC; Rev: CATCCAGGCTCTCTGGCTTCT).
